# A 23bp Indel Polymorphism in TLR2 Gene Enhances Inflammation and Disease Severity in Dengue

**DOI:** 10.1101/239988

**Authors:** A. Raj Kumar Patro, Sriprasad Mohanty, Aditya K. Panda, Birendra K. Prusty, Diwakar K. Singh, Sagar Gaikwad, Tanuja Saswat, Soma Chattopadhyay, Rina Tripathy, Bidyut Das, Balachandran Ravindran

## Abstract

**Background:** Dengue is the most rapidly spreading viral disease transmitted by the bite of infected *Aedes* mosquitos. Pathogenesis of dengue is still unclear; although host genetic factors, immune responses and virus serotypes have been proposed to contribute to disease severity. The development of high-throughput methods have allowed to scale up capabilities of identifying the key markers of inflammation. Since NS1 protein of dengue virus has been reported to activate immune cells towards enhanced inflammation through TLR2, we examined the role of a polymorphism, a 23bp deletion in 5’UTR region of TLR2 gene in patients with dengue (with and without warning signs) and correlated with plasma levels of inflammatory mediators with disease severity and viral serotypes.

**Methods:** Eighty nine patients classified as per WHO 2009 criteria during dengue outbreak in Odisha, India in 2016 were included in the current study. Presence of dengue virus (DENV) was demonstrated by detecting NS1 antigen, IgM capture ELISA and serotypes in circulation were discriminated by type-specific RT-PCR and/or sequencing. Sixty-one confirmed dengue cases were typed for TLR2 indel polymorphism and compared with 485 disease free controls. Plasma samples were assayed for 41-plex cytokine/ chemokines using Luminex bead based immunoassay.

**Results:** Presence of 23bp deletion allele of TLR2 gene was significantly more in patients with severe dengue in comparison to dengue fever cases (*p*= 0.03; Odds ratio 4.05) although the frequency of insertion (Ins) allele of TLR2 was comparable in healthy controls and dengue cases (82.4 and 87.9 % respectively). Seventy-three (82%) samples were found to be positive by NS1/IgM capture ELISA/ RT-PCR. DENV-2 was predominant (58%) during the outbreak. Among the host inflammatory biomarkers 9 molecules were significantly altered in dengue patients when compared to healthy controls. The increased levels of IFN-γ, GM-CSF, IL-10, IL-1Rα and MIP-1β correlated significantly with severe dengue.

**Conclusions:** The frequency of 23bp Indel mutation of TLR2 was comparable between healthy controls and dengue fever (with and without warning signs), suggesting that this indel mutation does not contribute significantly to susceptibility/ resistance to dengue; however, del allele of TLR2 gene was significantly more associated in patients with severe dengue symptoms when compared to dengue fever cases.

## INTRODUCTION

Dengue virus transmitted by *Aedes* mosquito, is an increasing global problem, with an estimate of 390 million infections per year and about 3.6 billion people at risk to dengue. India alone contributed to 34% of the total global infections (Bhatt *et al* 2013). Infection with the dengue virus (DENV) resulted with a spectrum of clinical manifestations ranging from asymptomatic infection, self-limiting, dengue fever (DF) with or without warning signs or to life threatening severe dengue (SD). Apparent dengue virus infections present with fever, headache, fatigue, nausea, chills, joint pain and dizziness. A few of the infections lead to life threatening severe dengue, manifest by vascular leakage results in shock, internal hemorrhage, organ impairment, results in death if untreated. The detail mechanisms of dengue pathogenesis lead to severe dengue is currently not clear.

There are four antigenically related but distinct serotypes of this virus, described as DENV-1, DENV-2, DENV-3, and DENV-4. The dengue virus is a positive strand RNA genome of 10.7 kb nucleotides, which encodes three structural (capsid, membrane and envelope) and seven non-structural (NS1, NS2A, NS2B, NS3, NS4A, NS4B and NS5) proteins. Infection with one of the four viral serotypes confers protective immunity against re-infection with the same serotype, while subsequent infection with different serotype results in severe dengue through antibody-dependent enhancement (ADE) result in cytokine storm (Halstead, 2007; Rothman et al, 2011; Screaton et al 2015).

On infection with dengue virus, initial innate immune response is elicited by production of inflammatory and antiviral molecules. Toll-like receptors (TLRs), a family of pattern recognition receptor plays important role in recognition of pathogen-associated molecular patterns (PAMPs). The non-structural protein-1 (NS1) protein of dengue virus induces pro-inflammatory cytokines through engagement of TLR2 (Chen et al. 2015). Several studies have shown the genetic variation in the TLRs and adapter molecules associated with disease susceptibility (Schwartz et al.2005); however, there is no information on the TLR2 23bp Indel in the 5’UTR downstream of NF-κB binding site for the disease severity in dengue.

An exacerbated host immune response marked by ADE plays an important role in development of severe dengue (Halstread et al., 2007; Guzman & Harris, 2016). Although expression of host soluble pro-inflammatory biomarkers proposed to be increased vascular permeability in endothelial cells in response to dengue infection have been reported in limited studies (Bozza et al., 2008; Rothman, 2011), the detail mechanisms involved in immune enhanced disease, hemorrhagic manifestations is not known. The availability of high throughput Luminex based cytokine bead assay allow potential correlation between genetic polymorphism of critical genes such as TLR2 and cytokine induction. Insights into biomarkers could also provide rational intervention strategies using cytokine therapy antagonists. Recent development of a dengue vaccine, Dengvaxia (WHO, 2017) has opened up avenues for reducing the disease burden. Because of the geographical preponderance of this virus, there are limited data on the circulating strains in India-more critically whether or not serotype specific host responses are associated with disease severity is not available. Since the vaccine is likely to be available in near future, it is imperative to examine the relative distribution of the circulating dengue serotypes.

We carried out the current study to examine the role of a hitherto unreported mutation, a 23bp deletion in 5’UTR region of TLR2 gene and a multiplex bead array for 41 host molecules and their correlation with disease severity. The study also demonstrates distribution of circulating serotypes in this geographical region and serotype specific immunity in severe dengue.

#### IMPORTANCE

Dengue is the most common arboviral infections observed worldwide with half of the global population are at risk. The increasing incidence of severe dengue is a major public health concern. An exacerbated host immune response marked via antibody-dependent enhancement (ADE) has been proposed to be crucial for dengue pathogenesis, while the role played by host genetic factors and viral determinants in dengue pathogenesis are unclear. Here we have demonstrated a correlation between TLR2 23bp indel mutation in 5’UTR region and serotype-specific host response in patients with severe dengue. Understanding the association of the Indel mutation in TLR2 gene in severe dengue, supported by the host inflammatory biomarkers and the dengue serotype may help guide in optimizing therapeutic efforts. Of the 41 cytokines examined five cytokines correlated significantly with disease severity. This study could be extrapolated to other flavivirus pathogenesis, since DENV NS1 shares a high degree of homology with all flavivirus NS1 proteins.

## METHODS

### Study population

From Jun 2016 through Dec 2016, 89 subjects presented at the S.C.B. Medical College & Hospital, Cuttack, Odisha, India, were enrolled in the study. The subjects originated from three different states; Odisha, West Bengal and Jharkhand, of the eastern region of India. Subjects with high fever, nausea, joint pain, dizziness, shock-like state, rash, drop in blood pressure and amplification of primary symptoms, vascular system damage and bleeding, consistent with WHO criteria 2009 (Alexander et al., 2011; Katzelnick et al., 2017), were included in the study. Blood samples were collected and immediately transported in ice to the laboratory. The samples were aliquoted and immediately processed for the experiment and rest kept at −80^0^C for further use. Healthy control subjects (485, mostly institutional volunteers and disease free persons accompanying patients in S.C.B. Medical College, Cuttack, India, were included in the study. Institutional Human Ethic Committee Review Board approval was obtained at each study site, from Institute Human Ethics Review Committee of S.C.B. Medical College and Hospital and Institute of Life Sciences, India, and blood samples were collected after informed written/oral consent was taken.

### Serology

Detection of the dengue NS1 antigen and IgG on serum samples were done using BeneSphera Dengue Test Kit (Avantor, PA, USA) according to manufacturer’s instructions. Qualitative detection of IgM antibodies to dengue antigens for recent infection in serum samples were performed using the Panbio® Dengue IgM Capture ELISA kit (Panbio, MA, USA).

### Detection of dengue virus and typing

RNA was extracted from the serum samples using commercial viral RNA extraction kit (QIAamp viral RNA extraction kit, Qiagen, Germany) and immediately converted to cDNA using high capacity RNA to cDNA conversion kit following manufacturer’s instructions (ABI-Applied Biosystems, USA). The cDNA was subjected to polymerase chain reaction amplifying 511 bp sequences spanning the dengue virus capsid premembrane region for detection of dengue. Further, type specific primers used to discriminate individual serotypes as reported previously (Lanciotti et al., 1992, Saswat et al., 2015). The PCR products were resolved on 2% agarose gel and visualized with staining ethidium bromide. Selected amplicons were gel purified and sequenced to ascertain the findings using ABI genetic Analyzer and Big Dye Terminator DNA sequencing kit (Applied Biosystems, Foster City, CA, USA) at the Institute of Life Sciences, core sequencing facility. The sequences were submitted to national centre for biotechnology information (NCBI).

### Detection of TLR2 23bp Indel Mutation

DNA was extracted from the 485 healthy controls and 61 dengue positive subjects from peripheral blood specimens using a commercial spin column kit (QIAamp DNA extraction kit, Qiagen, Germany). The extracted DNA subjected to polymerase chain reaction using primers spanning the TLR2 23bp indel region with forward-sense 5’-CACGGAGGCAGCGAGAAA-3’ primer, and reverse 5’-CTGGGCCGTGCAAAGAAG-3’ primer as reported earlier (Panda et al., 2017). After initial denaturation for 4 minutes at 95^0^C, the samples underwent 35 cycles of denaturation 95°C for 30s, annealing at 60°C for 30s, and extension 72°C for 30s with a final extension at 72^0^C for 7 min. PCR products were resolved on a 2% TAE agarose gel and visualized using staining with ethidium bromide and image captures using gel documentation system. Amplicons with a single band at 286 bp was classified as wild type (ins/ins), a single 263bp band as homozygous mutation (del/del), and products displaying two bands of 286bp and 263bp were treated as heterozygous mutation (ins/del). A second set of primers encompassing the Indel region was designed using the Primer 3 software (Untergasser et al., 2012) and used in PCR-Sequencing to detect the mutation. The sequencing primer for Forward Sense: 5’ CCACCTGCCTGGAACTCA 3’ and anti-sense: 5’ GTGTAGCCCCTGCTTCTCC 3’ amplifying 508 bp product. The 23bp deletion region in 5’ UTR of TLR2 gene sequence is 5’ CAGGCGGCTGCTCGGCGTTCTCT3’ (Figure 6). Selected PCR products were column purified (using Qiagen PCR purification kit, Qiagen, Germany) and sequenced using ABI genetic Analyzer and Big Dye Terminator DNA sequencing kit (Applied Biosystems, Foster City, CA, USA) at the Institute of Life Sciences, core sequencing facility. Samples were genotyped by different laboratory personnel for ensuring reproducibility of typing.

### Analyses of 41plex cytokines using multiplex Luminex system

The levels of 41 cytokine molecules were measured using Luminex based bead array the MILLIPLEX® MAP Human Cytokine / Chemokine panel on LuminexxMAP® platform using a magnetic bead format (MILLIPLEX® Analytes, Millipore, MA, USA,) for the following biomarkers: EGF, FGF, Eotaxin, TGFα, G-CSF, Flt, GM-CSF, Fractalkine, IFNα2, IFNγ, GRO, IL-10, MCP, IL-12P40, MDC, IL-12P70, IL-13, IL-15, sCD40L, IL-17A, IL-1RA, IL-1α, IL-9, IL-1β, IL-2, IL-3, IL-4, IL-5, IL-6, IL-7, IL-8, IP-10, MCP-1, MIP-1α, MIP-1β, TNFα, TNFβ, VEGF, PDGF-AA, PDGFAB-BB and RANTES. In brief, plasma samples were tested for the levels of the cytokines, using the multiplex immunoassay containing fluorescent labeled beads conjugated with specific monoclonal antibody for the target molecule, according to the manufacturer’s recommendations. Samples were considered for analysis using the standards that were run in each of the plates. Each run contained appropriate quality controls and results were interpreted from the standard run in duplicates using Bioplex 200 system (Bio-Rad).

### Venn diagram

Relationship between circulating dengue serotypes was made by Venn diagram using the online versed Venny 2.1 software. [http://bioinfogp.cnb.csic.es/tools/venny/].

### Principal component analysis (PCA) and Partial Least Squares Discriminant Analysis (PLS-DA)

MetaboAnalyst 3.0 software (http://www.metaboanalyst.ca) for multivariate Principal Component Assay (PCA) and Partial Least Squares Discriminant Analysis (PLS-DA) was used to comprehensively analyze the 41-plex cytokine bead array. Analysis of the 41-plex cytokine expression for different groups, i.e., TLR2 23bp Indel in dengue serotype, control vs dengue fever and dengue fever versus severe dengue by high dimensional spectral features using multivariate PCA and further discrimination of inter-class variance between the groups were made by the PLS-DA [Xia et al., 2016]. The variable importance in the projection (VIP) score for the multiplex cytokine profiles in PLS-DA that summarizes contributions of the variables gets depicted in the model. The cytokine with highest VIP scores and hence most contributing variable and the least discriminating parameter the in PLS-DA model were depicted by color code.

### Statistical analysis

Statistical analyses were performed using software GraphPad Prism® software PRISM 6 version, GraphPad Software, Inc., CA, USA. The cytokines/ chemokine data were analyzed using Student’s t-test Student’s test or ANOVA as appropriate. Fisher’s exact test was used for comparison of genotype, allele frequencies and to test association of combined genotype distribution among various clinical categories. Proportions were computed with 95% confidence intervals (95% C.I.) according to the Poisson distribution and a ‘p’ value less than 0.05 was considered as statistically significant.

## RESULTS

### Population demographics

The demographic characteristics were as follows: the age distribution range from 17 to 72 years, mean age 33.2 ± 12.5 (SD) years. Males constituted 77% while the rest were females. When stratified for age, majority of the subjects were in the age group of in 25–34 years age group. Detailed clinical presentations in dengue and severe dengue are shown, Table S1 and Table S2. Among the clinical manifestations of retro-orbital pain, itching and bleeding manifestations were statistically significant between the dengue fever (with and without warning signs) and severe dengue group (Table S1). Based on the clinical conditions, subjects were treated with conservative fluid management, prophylactic antibiotic treatments and considered for platelet transfusion based on the disease severity as per the WHO guidelines.

### 23bp Indel at 5’UTR of TLR 2 Gene distribution in Dengue Patients

Among the 485 healthy controls, the ins/ins genotype was found in 69.7% (338/485), ins/del heterozygocity in 25.5% (124/485) and del/del homozygocity in 4.7% (23/485) subjects (Table 1). Among the 61 dengue subjects examined (54 with dengue fever and 7 with severe dengue), the following were the distribution: wild type in 83.3%, heterozygotes in 9.25% (5/54), while del/del homozygotes in 7.4%; details are presented in Table 2. The difference between healthy controls and mild dengue cases were comparable, while the difference between the mild dengue and severe dengue is significantly different as shown in Table 2 (p= 0.03, OR = 4.05; 95% C.I-1.17-13.9).

**Table 1.** Distribution of TLR2 23bp [Ins/Del] genotype & allele in controls (healthy subjects) vs dengue fever (with & without warning signs)

| **Genotype and allele distribution of TLR2 23bp (ins/del) polymorphism in controls vs dengue subjects** |  |  |  |
| --- | --- | --- | --- |
| **Genotype** | **Healthy controls**<br>**(n = 485)** | **Dengue Fever**<br>**(n = 54)** |  |
| ins/ins | 338 (69.7%) | 45 (83.3%) |  |
| ins/del | 124 (25.5%) | 5 (9.25%) |  |
| del/del | 23 (4.7%) | 4 (7.4%) |  |
| **Allele** | **Healthy controls**<br>**(n = 970)** | **Dengue Fever**<br>**(n = 108)** |  |
| ins | 800 (82.4%) | 95 (87.96%) |  |
| del | 170 (17.5%) | 13 (12.04%) |  |
| **Genotype** | **P-value** | **Odds ratio** | **95% C.I.** |
| ins/ins vs. ins/del | 0.0093 | 0.302 | 0.1175-0.7805 |
| ins/ins vs. del/del | 0.5483 | 1.306 | 0.432- 3.9495 |
| **Allele** | **P-value** | **Odds ratio** | **95% C.I.** |
| ins vs del | 0.1766 | 0.644 | 0.3524- 1.1766 |

\*p value (Fisher's exact test, two tailed)
Association of TLR2 23bp ins/del genotype with severe dengue computed through Fisher's exact test. Statistical analyses were performed using software GraphPad Prism® software PRISM 6 version, GraphPad Software, Inc., CA, USA.
Fisher's exact test, two tailed was used to compare proportions. Proportions were computed with the corresponding 95% confidence intervals (95% C.I.) according to the Poisson distribution. A 'P' value less than 0.05 was considered as statistically significant.

**Table 2.**
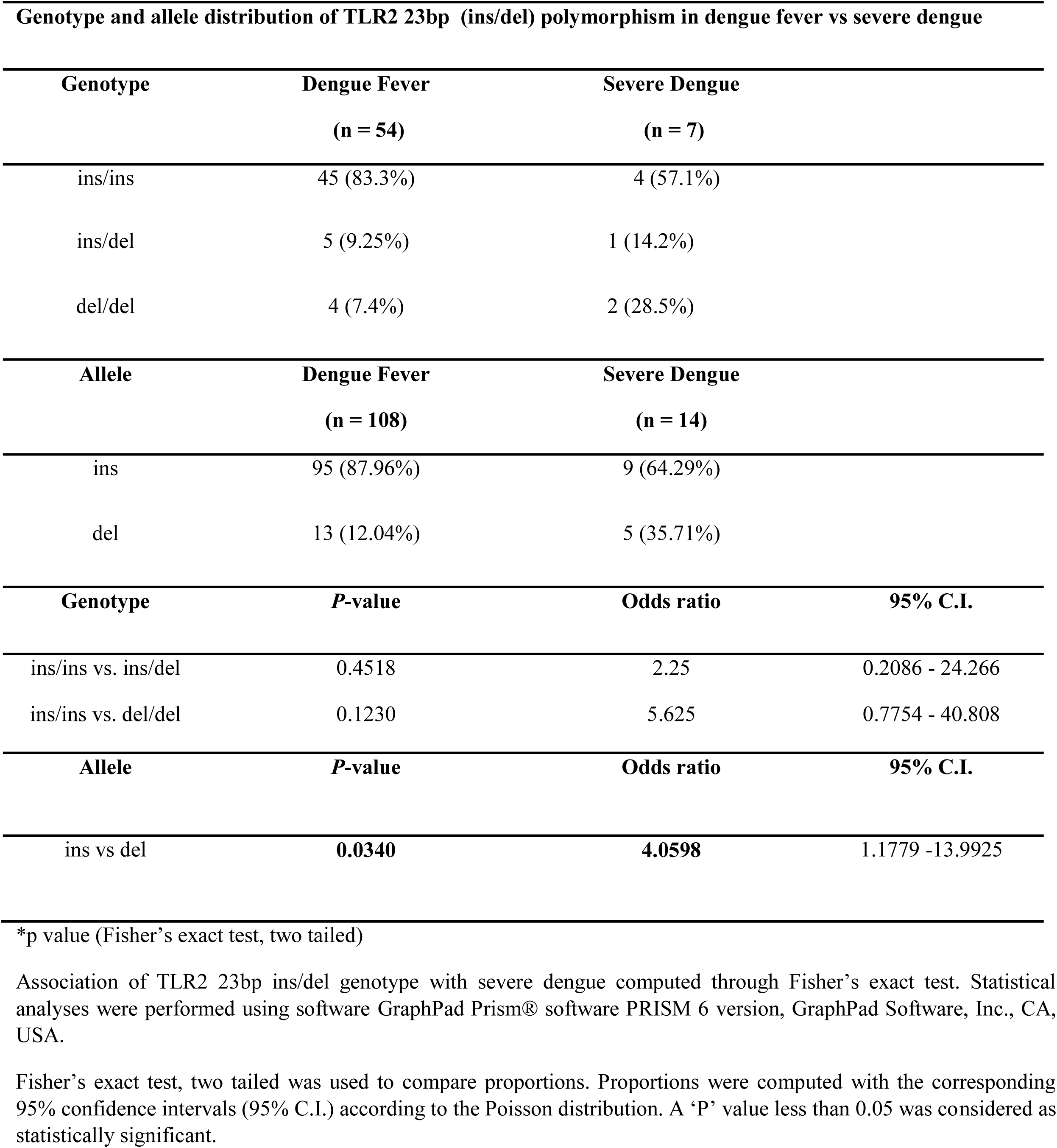
Distribution of TLR2 23bp [ins/del] genotype and allele in dengue fever (with & without warning signs) vs severe dengue

### Multiplex Cytokine Bead Array (Inflammatory markers as predictors of the dengue severity)

In dengue patients the following plasma cytokine levels were significantly elevated when compared to healthy controls: of GM-CSF, IFNγ, IL-10, 1L-15, IL-8,MCP1,IL6,MIP1β and TNFα levels. Increased plasma cytokine levels of IFNγ, GM-CSF, IL-10, IL1Rα and MIP-1β significantly correlated with severe dengue (Figure 2). Further, the IFN-γ and IL-10 levels were significantly increased in the TLR2 23bp del allele as compared with the 23bp ins allele (Figure 3). In contrast, RANTES and sCD40L were significantly elevated in the 23bp ins allele as compared to the 23bp del allele (Figure 4).

**Fig. 1.**
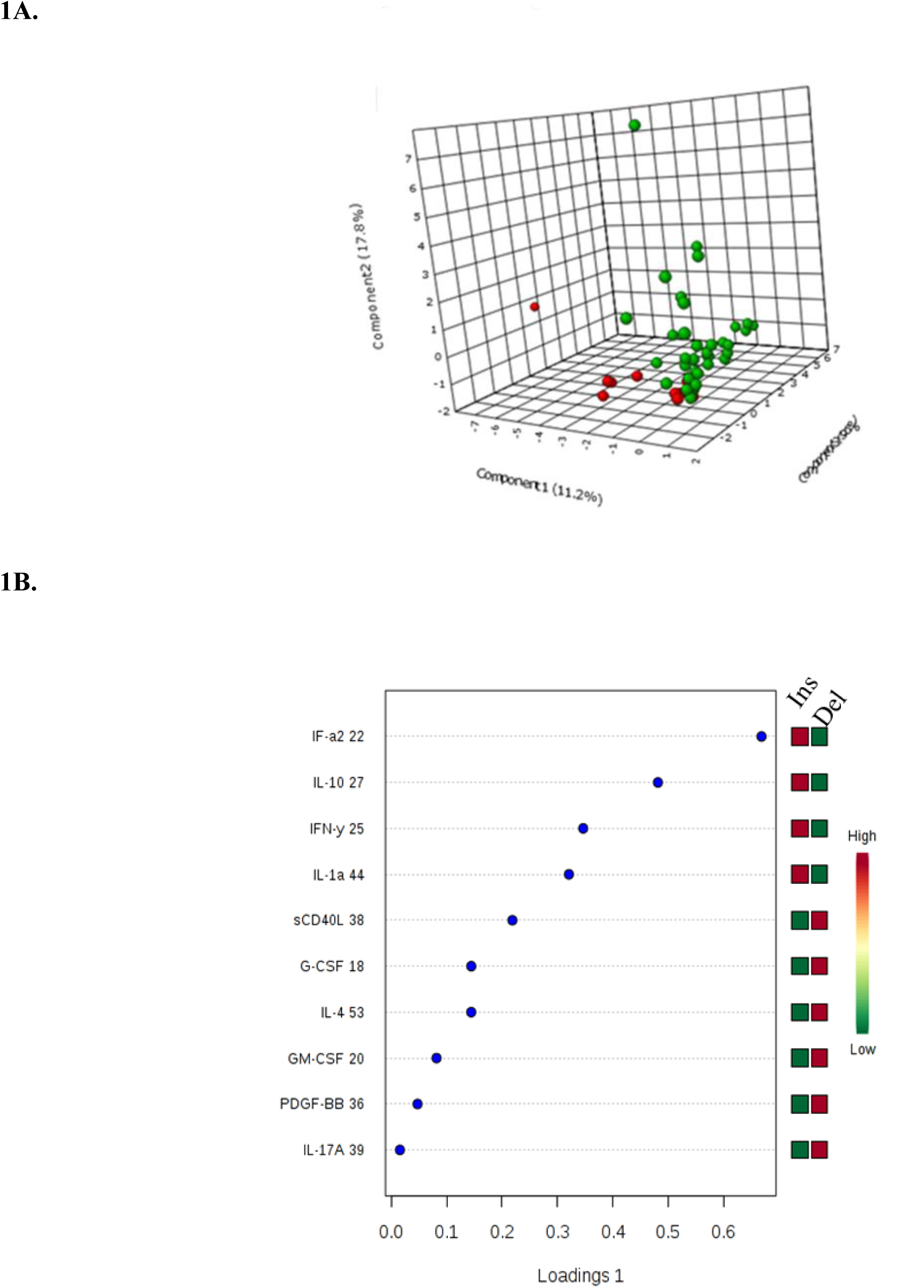
Partial least square discriminant analysis (PLS-DA) & variable importance in projection (VIP) scores of TLR2 Ins vs Del allele & Cytokine profile in Dengue Subjects. Principal component analysis (PCA) and Partial Least Squares Discriminant Analysis (PLS-DA) depicting the cluster between the ins allele and the del allele subjects in their cytokine expression profile (1A) The red color indicates for the del allele, while the green color depicted for the ins allele. The variable importance in the projection (VIP) score for the multiplex cytokine profile summarizes the contributions of the variables depicted in the model (1B).

**Figure 2.**
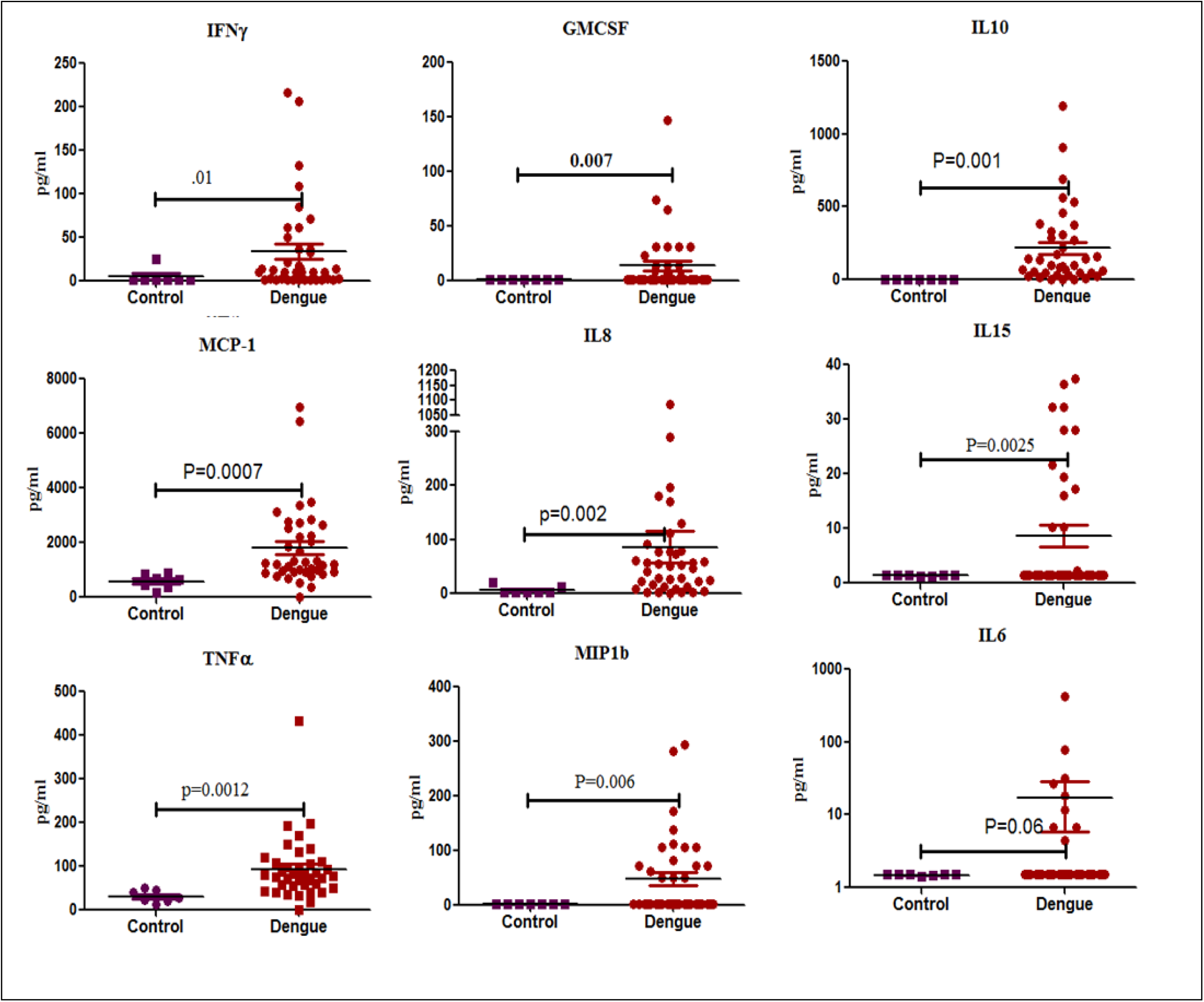
Cytokine levels in healthy controls and dengue positive subjects. Scatter plot showing the cytokine expression profile among the control vs dengue positive subjects. Of the 41 cytokine examined through Luminex bead array only nine cytokines found to be associated with the dengue positivity when compared to the controls. Statistical analyses were performed using software GraphPad Prism® software PRISM 6 version, GraphPad Software, Inc., CA, USA. Proportions were computed with the corresponding 95% confidence intervals (95% C.I.) according to the Poisson distribution. A ‘P’ value less than 0.05 was considered as statistically significant.

**Figure 3.**
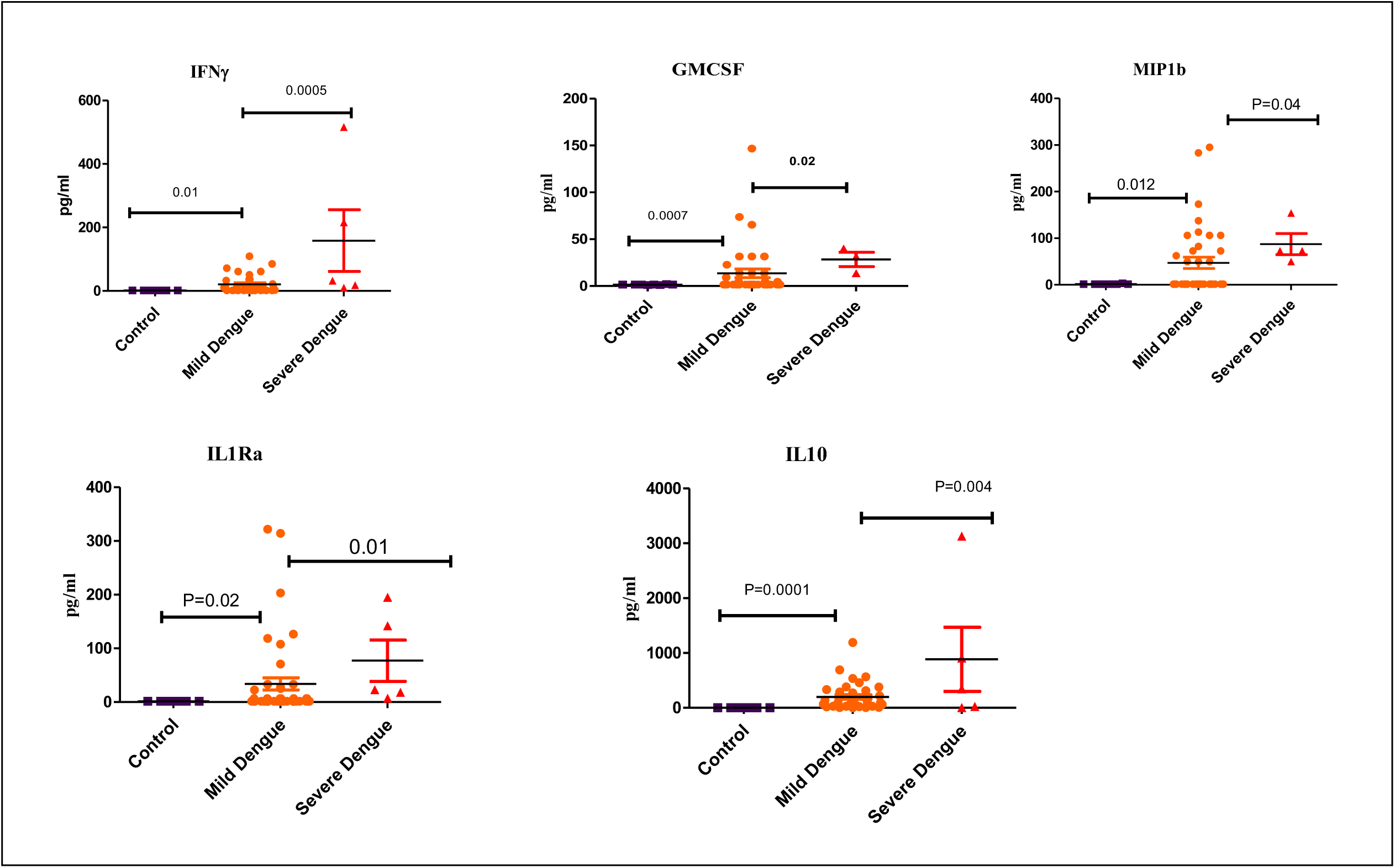
Cytokine levels in control, dengue fever (with & without warning signs) and severe dengue. Scatter plot showing the cytokine expression profile among the control, dengue (with & without warning signs) and severe dengue subjects. Of the 41 cytokine examined through Luminex bead array, five cytokines found to be associated with disease severity (SD), when compared to the dengue positive (with & without warning signs) and the controls. Statistical analyses were performed using software GraphPad Prism® software PRISM 6 version, GraphPad Software, Inc., CA, USA. Proportions were computed with the corresponding 95% confidence intervals (95% C.I.) according to the Poisson distribution. A ‘P’ value less than 0.05 was considered as statistically significant. The cytokines IFN-γ, GM-CSF, IL-10, MIP-1β and IL1Ra could be the predictor disease severity.

**Figure 4.**
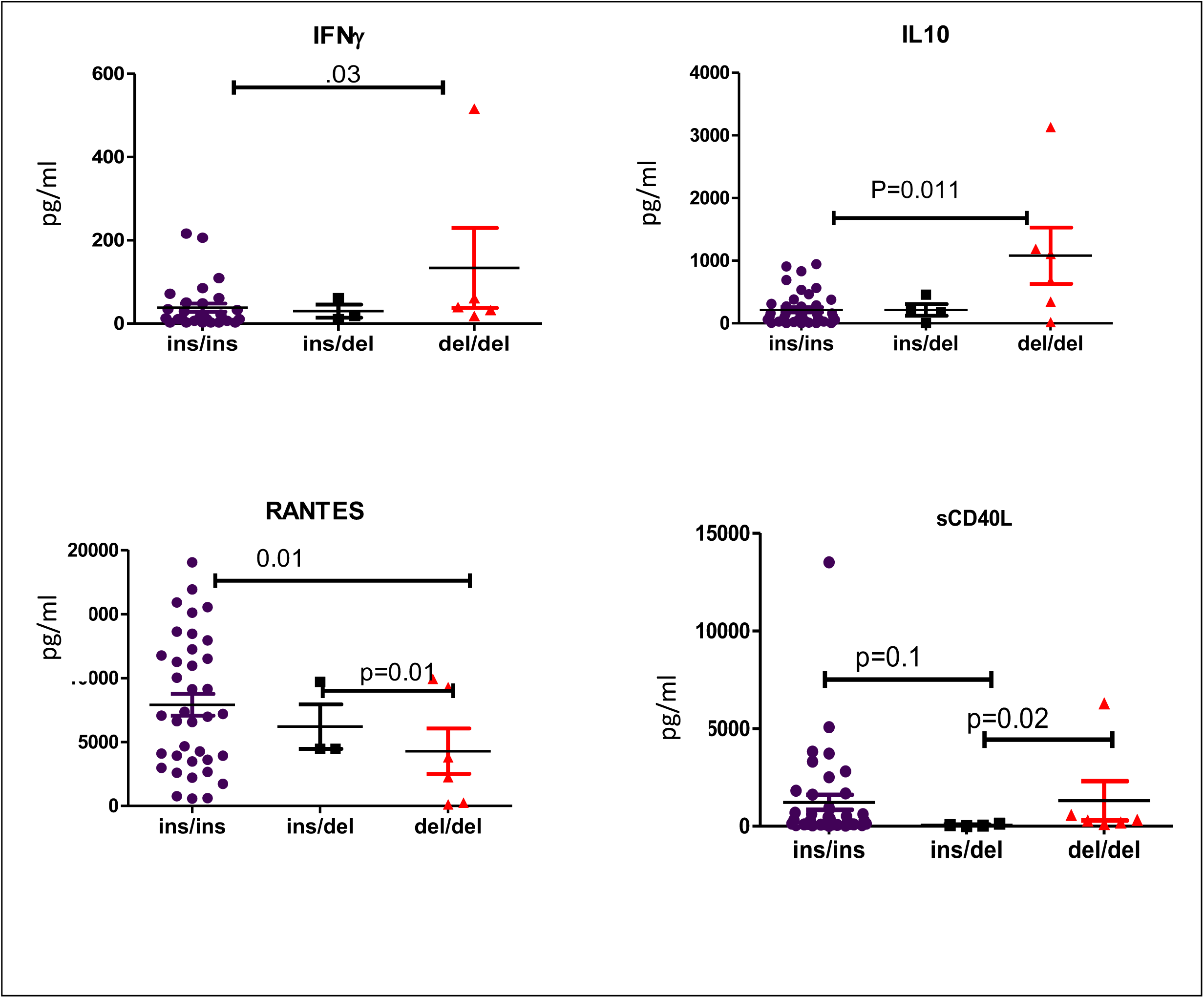
Cytokine levels in subjects with TLR2 23bp ins allele and del allele. Scatter plot showing the cytokine expression profile among the dengue positive subjects with ins and del allele. Of the 41 cytokine examined through Luminex bead array, cytokines IFN-γ and IL10 significantly elevated in the severe dengue subjects. However, the molecules RANTES and the sCD40L which are derivatives of platelets inversely correlated in disease severity. Statistical analyses were performed using software GraphPad Prism® software PRISM 6 version, GraphPad Software, Inc., CA, USA. Proportions were computed with the corresponding 95% confidence intervals (95% C.I.) according to the Poisson distribution. A ‘P’ value less than 0.05 was considered as statistically significant.

**Figure 5.**
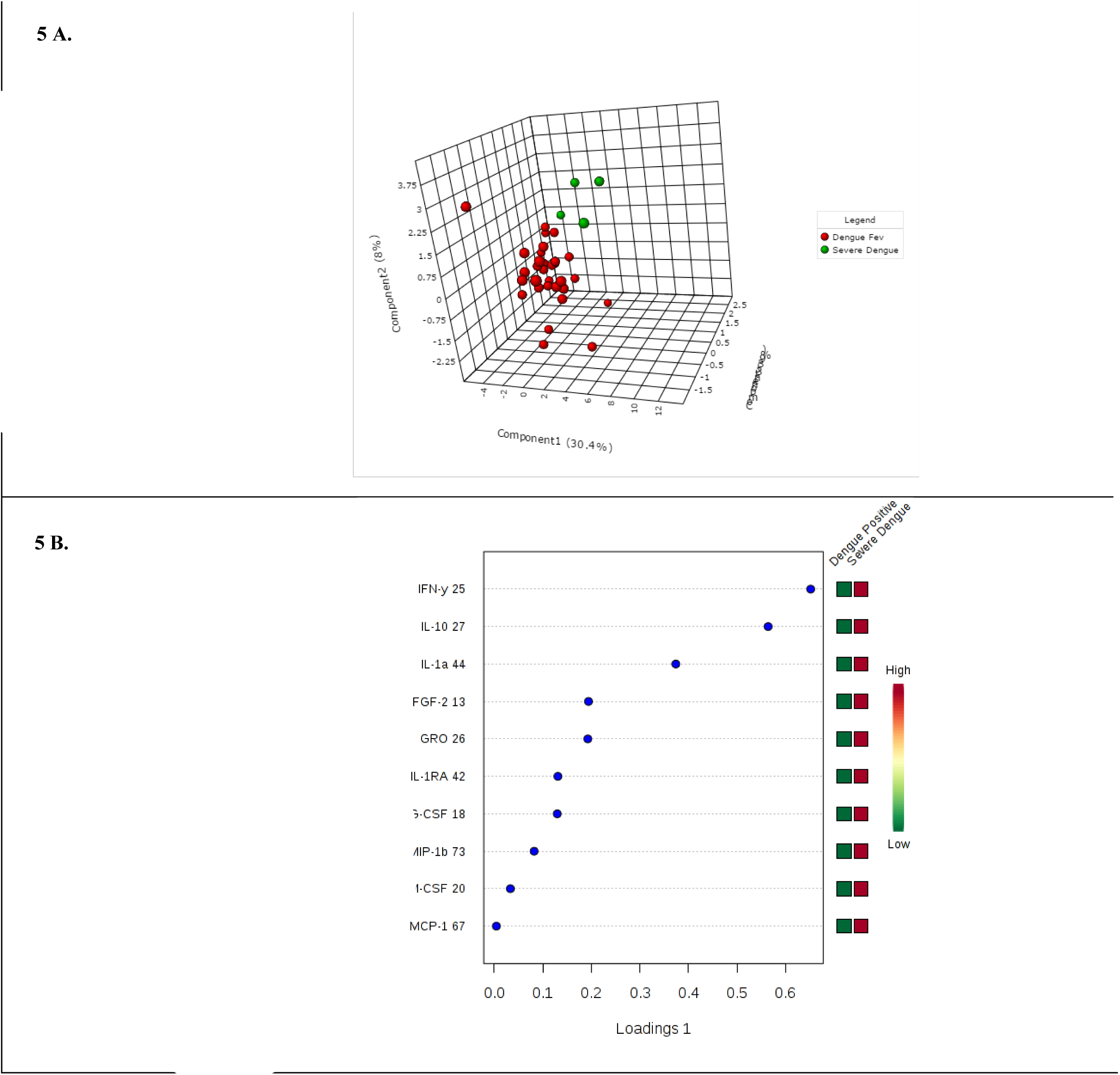
Principal component analysis (PCA) of dengue positive patients: dengue (with & without warning signs) and severe dengue. Principal component analysis (PCA) and Partial Least Squares Discriminant Analysis (PLS-DA) A depicting the cluster between the multiplex cytokine expression profile between the dengue fever (with & without warning signs) [DF] and the severe dengue [SD] subjects. The red circle represents for the dengue fever [DF], while the green color depicted for severe dengue [SD] subjects cluster (5 A). The variable importance in the projection (VIP) score for the multiplex cytokine profile summarizes the contributions of the variables depicted in the model (5B). Both the dengue and severe dengue group clustered distinctly in terms of the expression profile of the soluble biomarkers

**Figure 6.**
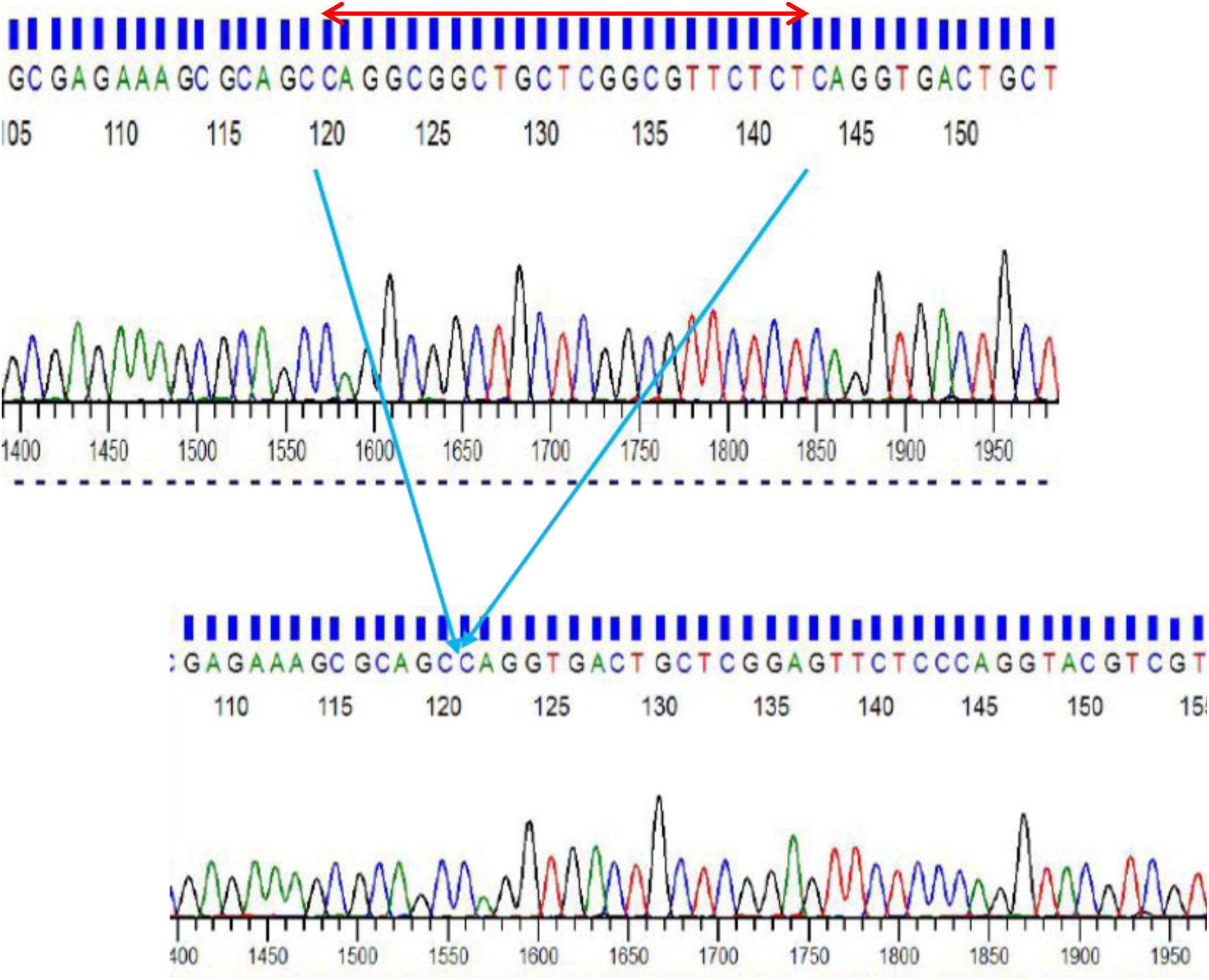
Chromatograms image of part of TLR2 23bp Indel sequence showing site of mutation. Chromatogram showing ins/ins (Top) and bottom showing del/del (bottom) region Deletion mutation region: CAGGCGGCTGCTCGGCGTTCTCT

### Dengue Testing

Of the 89 subjects evaluated for the study (73) 82% were found positive by NS1/ IgM capture ELISA/ RT-PCR. Out of these 73 patients 90.41% presented with dengue fever (with and without warning signs) and 9.58% patients were categorized as severe dengue. Of the positive samples, NS1 antigen test detected in 60.3% subjects (44/73), while the IgM capture ELISA detected in 80.82% (59/73) for IgM recent infections. Dengue serotype-specific RT-PCR discriminated for DENV in 36 (49.3%) [36/73] 95% C.I: 38.17%-60.53%] patients; single infection for one serotype was observed in 61.11% while mixed infection with more than one serotype were detected in 38.88% patients. Overall there were 50 infections in 36 RT-PCR positive subjects for the four dengue serotypes DENV-1 in 24%, DENV-3 in 8%, DENV-4 in 10% while majority (58%) were positive for DENV-2 either as single or as multiple infection with other serotypes (Table 3). Further, selected samples were sequenced to ascertain the findings.

**Table 3.**
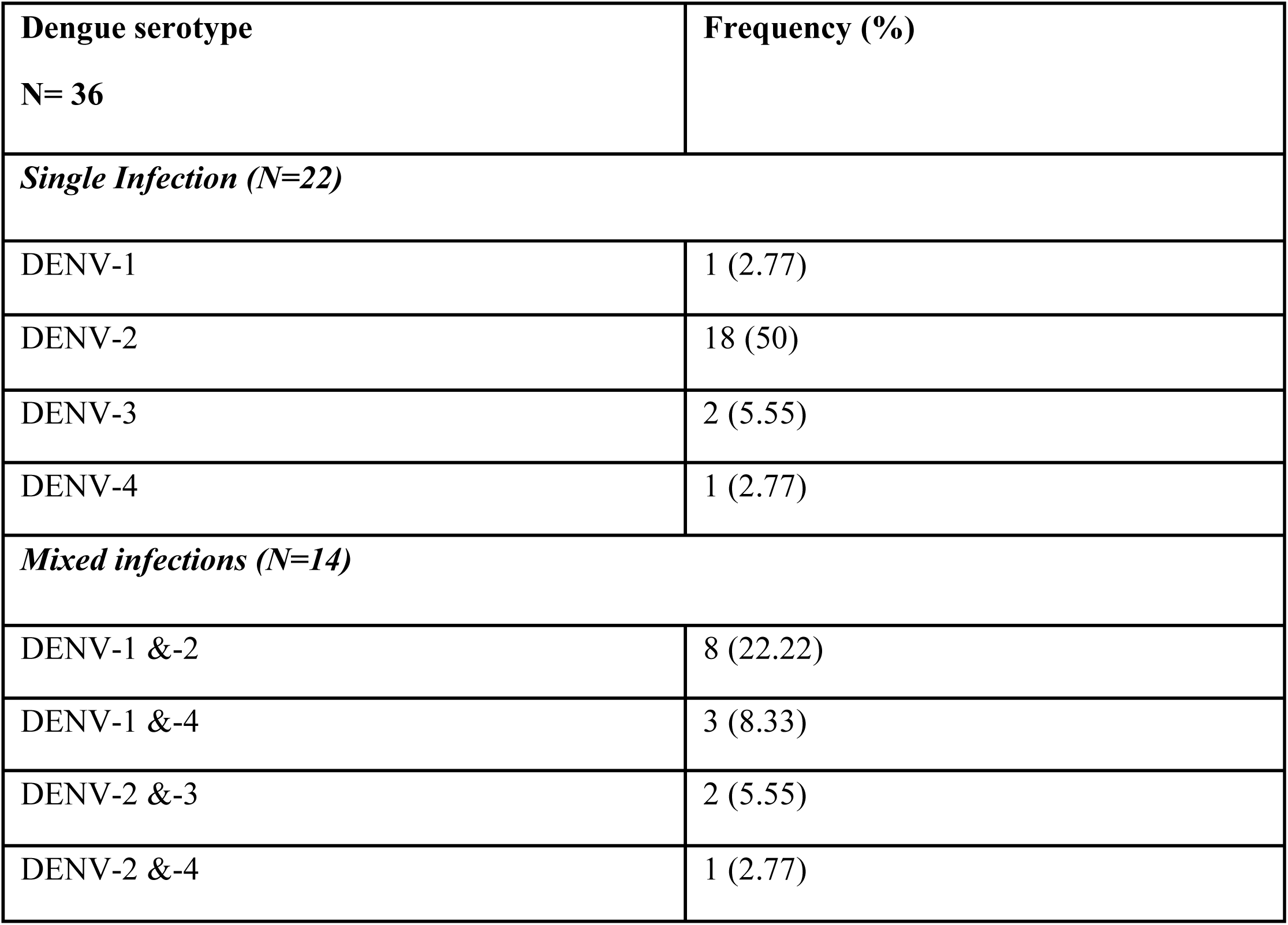
Circulation of dengue serotypes in 2016 outbreak in eastern India. Typing of dengue positive subjects by type specific RT-PCR (N=36). Single infection seen in 22 subjects, while mixed infections were seen in 14 subjects.Figure 5. Principal component analysis (PCA) of dengue positive patients: dengue (with & without warning signs) and severe dengue

### Principal Component Assay & Venn diagram

Multivariate analysis by Principal Component Assay for the 41 cytokine expression revealed two distinct clusters for the ins and del allele and the PLS-DA & VIP score of TLR2 ins/ins vs del/del allele & Cytokine profile in dengue subjects clearly distinguished the two groups (Figure 1). When cytokine expression profiles were compared between healthy controls and dengue positive patients with single and mixed infections grouped in two separate clusters (Figure S1). Further the cluster differentiated inflammatory molecules between dengue fever and severe dengue (Figure S2). PLS-DA and VIP score also effectively differentiated DENV-2 from DENV -1, -3, -4 (pooled as a single group) for the biomarkers tested in this study (Figures S3).

### Venn diagram

Comparison and interrelationship between the circulating dengue serotypes in mild dengue vs severe dengue were analyzed using Venn diagram and the results are shown in (Figure S4). We found mixed infections of virus subtypes were associated in dengue fever viz., DENV-1 and DENV-2 (18.8 %), DENV-1 and DENV-4 (6.3%). In addition, severe dengue was associated with mixed infections of DENV-2 and DENV-3 (20%) subjects.

### Sequences and NCBI accession number

The sequences of circulating dengue strains submitted to NCBI (Accession number-MF069161, MF441493).

## DISCUSSION

One of the unsolved conundrum in Dengue pathogenesis is the consistent observation that only a small proportion of dengue infected patients present with severe clinical manifestations. The mechanism of development of the severe dengue in a small cohort of patients has been a subject of intense study in recent years. Host genetic factors and difference in immune responses in patients have been proposed as possible reasons (Rothman et al., 2011) The dengue virus NS1 has been demonstrated to play a critical role in inducing vascular damage causing dengue pathology (Beatty et al., 2015; Puerta-Guardo et al., 2016; Glasner et al., 2017). In the context reports on NS1 being a TLR2 ligand and activating innate immune cells leading to hyper-inflammation observed in severe dengue (Chen et al., 2015), in this study we studied the Indel polymorphism of a 23bp sequence in 5’ UTR region of TLR2 in humans. We had shown that this unique primate specific polymorphism absent in rodents contributes to TLR2 mediated inflammation in Malaria and Sepsis (Panda et al., 2017). Activation of TLR2 is known to promote production of type I interferon, one of the key components of anti-viral immunity. In this context, we examined the role of a 23bp deletion in 5’UTR region of TLR2 gene genomic variant 60bp downstream of NF-κB binding site (Johnson et al., 2007) in dengue and severe dengue cases. While the frequency of Indel polymorphism between controls and mild dengue cases was comparable (suggesting that this polymorphism) does not contribute significantly to increased susceptibility/resistance to dengue, a statistically significant difference (p <0.03; Odds ratio 4.05) was observed between mild versus severe cases of Dengue. These observations suggest that patients with del/del genotype are more prone to develop severe dengue if they are infected with dengue virus. We had reported earlier that the del/del genotype is restricted to lower primates, while Chimpanzee, Orangutan, Neanderthal and Human harbored 23bp nucleotide insertion that appears to have offered evolutionary advantage against adverse disease outcome in Malaria and Sepsis (Panda et al., 2017). The results of this study strengthen this scenario in dengue infections also. Our current observations that del/del polymorphism is associated with elevated inflammatory cytokine levels in dengue patients further consolidates our view and suggests ins allele confers an evolutionary advantage to the host in dengue also. DENV is known to infect various immune cells including monocytes and dendritic cells, which majorly contributes to production of inflammatory and/or antiviral cytokines. While unregulated inflammation leads to exacerbated pathogenesis and tissue/organ injuries followed by death (Costa et al., 2013).

Patients with dengue present a ‘cytokine storm’ accompanied by excessive secretion of cytokines and chemokines. Therefore, analysis of cytokine and chemokine levels could be a tool for predicting status/severity of the disease. The multiplex bead array used in our study revealed the cytokines GM-CSF, IFN-γ, IL-10, IL-15, IL-8, MCP-1, IL-6, MIP-1β and TNF-α levels were significantly higher in dengue subjects compared to healthy controls. Other studies have reported increased significantly elevated plasma levels of interferon-γ and IL-10 levels in severe dengue (Green et al 1999, Libraty et al., 2002, Nguyen et al., 2004; Bozza et al., 2008, Priyadarshini et al., 2010). A recent report from GWAS studies on host on asymptomatic and clinical dengue subjects reveled most of the changes seen in expression profile belongs to immune responses (Simon-Lorière et al., 2017). However this study examined the transcriptome profile (rather than plasma levels) in patients with and without symptomatic dengue. In our study, levels of IL-15 and TNF-α were significantly more in dengue patients when compared with the controls. Further in the current study all patients with all four serotypes were analyzed while Simon-Lorière et al., 2017 had analyzed patients with only serotype 1 Dengue cases. The plasma cytokine levels of IFN-γ, GM-CSF, IL-10, MIP-1β and MCP-1 were significantly elevated (p= 0.0005, 0.02, 0.0004, 0.04, 0.01 respectively) in severe dengue patients in our cohort as compared to dengue fever cases. Expression of IFN-γ and IL10 increased in del allele compared to ins allele, while RANTES and the sCD40L has decreased in the del allele in compare to the ins allele (Figure 4). In dengue infection, sCD40L and RANTES are released by platelets, which play an important role in immune activation of antigen presenting cells thereby enhances IFNγ production and hence, the decrease in the RANTES could be due to thrombocytopenia in these subjects.

In this context, we observed that sCD40L and RANTES are decreased in the 23bp TLR2 del allele genotype which correlates with the disease severity. In concurrence of the findings, studies have shown the decreased level of sCD40 and RANTES in severe dengue (Soo et al., 2017). Thus levels of these plasma cytokines could be used as biomarkers of both severity and possibly for prognosis and their antagonists could be potentially be used as therapeutic agents. This is the first study to demonstrate the involvement of the 23bp TLR2 Indel mutation and DENV serotype-2 association with disease severity accompanied by elevated plasma inflammatory mediators. This study thus underscores the interplay of serotype-specific immune response and TLR2 Indel polymorphism in determination of dengue disease severity. Similar report on the disease severity by the dengue virus serotype-2 was demonstrated in Nicaragua with serotype specific immunity (Rico-Hesse et al., 2007; OhAinle et al., 20111)

The data presented in this communication however has limitations, since the numbers of cases with severe dengue were not high during this outbreak. However, it is unlikely to have any effect on the broad findings of the key molecules in terms of the data outcome. Adequate age and sex matched controls were included for the detection of the Indel polymorphism. Besides conducting luminex assay for 41 host molecules in circulation, all samples were analyzed by multiple methods for diagnosis and through consensus and type-specific RT-PCR.

Screening a much larger cohort of patients may have contributed to higher stringency for analysis of frequency of each of the four viral serotypes and levels of inflammatory molecules and distribution of 23bp Indel polymorphism of TLR2 gene in Dengue patients. Furthermore, the findings of this study could also be valuable for other flavivirus infections, since the TLR2 mediated inflammation due to DENV NS1 protein, which shares high degree of homology with all flavivirus NS1 protein (LindenBach et al Fields 2013).

In conclusion, this study provides first evidence of association of the TLR2 23bp indel polymorphism associated with enhanced inflammation and determines Dengue Disease Severity.

## Supplementary Data

Additional data available in Supplement section

**Notes**

## Acknowledgements

The contribution of Dr. Naga Jogoyya and Pulak Mohanty in sequencing of the samples at the Institute core facility is gratefully acknowledged. We also thank Subrata Mohanty and Aisurya Ray for collection and processing of the samples.

## Disclaimer

The funders had no role in the study design, data collection and analysis, preparation of the manuscript and decision to publish.

### Funding

A. Raj Kumar Patro, was supported by start-up grant from Department of Science and Technology, Science Engineering Research Board (Grant no. YSS2015/00389), Govt. of India. The Institute of Life Sciences, Bhubaneswar is supported by core funds made available by Department of Biotechnology, Government of India.

